# Using spatially-explicit plant competition models to optimise crop productivity in intercropped systems

**DOI:** 10.1101/2021.05.04.442600

**Authors:** Laura Stefan, Nadine Engbersen, Christian Schöb

## Abstract

Intercropping, by capitalizing on positive biodiversity–productivity relationships, represents a promising option to increase agricultural sustainability. However, the complexity and context-dependency of plant–plant interactions can make it challenging for farmers to find suitable crop combinations. Furthermore, intercropping is usually implemented with standard inter-row spacing and plant densities based on monoculture practices, which might not be the ideal configuration to maximise yield. Therefore, here we present a spatially-explicit method based on plant ecological interaction models that allowed to optimize crop species combinations and spatial configurations for maximal yield in intercropped systems. We tested this method with three crop species, namely oat, lupin, and camelina. In a first step, field experiments in which crop density was varied provided us with indications on which species would compete more with each other. The results showed us that oat and camelina strongly competed with each other. In addition, the distance experiments allowed us to understand how the changes in yield associated with the presence of neighbours vary with distance. This allowed us to find the sets of parameters (identity of neighbours, sowing density, distances between individuals) that optimises intercrop yield (measured as Land Equivalent Ratio [LER]) for the three considered species. Specifically, we show that alternating rows of species led to higher LERs than a homogeneous species mixing. In addition, for each spatial configuration considered, we provide indications for the optimal inter- and intra-row distances and information about relative yield losses with suboptimal planting patterns. By modelling crop yield from simple and reproducible density and distance experiments, our results allow to optimize crop mixtures in terms of species combinations and spatial configurations.

## 1. Introduction

Intercropping is the agricultural practice of growing more than two crop species on the same field during a significant part of their life cycle (Vandermeer 1989). By utilizing positive biodiversity-ecosystem functioning relationships, crop mixtures have been recognized as a promising way to make agriculture more sustainable (Tamburini et al. 2020).

In monospecific crop stands, sowing densities and intra- and inter-row distances are generally optimized for maximal population performance at the field scale (Weiner 2017). These parameters do not necessarily maximize individual performance, as there is a trade-off between optimal individual performance and optimal population performance in monocrops (Anten and Vermeulen 2016). Intercropped systems add a layer of complexity to this problem, as the goal might not only be to maximize population level yield, but also community (i.e. multi-species) level yield. Moreover, in monocultures, optimal sowing densities and intra- and interrow distances generally vary among species (Olsen et al. 2006, Testa et al. 2016). These elements suggest that optimal spacing might vary in an intercrop compared to a monocrop. However, there is little research on spatial configurations in intercrops, particularly when more than two species are considered. For instance, it remains unclear whether it would be more efficient for community-level yield to mix species as much as possible rather than keeping them in lines or strips (Galanopoulou et al. 2019, Cheriere et al. 2020).

The ecological theory of coexistence suggests that interactions between species, and competitive interactions in particular, are core drivers of individual-, population- and community-level fitness (Chesson 2000). Typically, individual-based growth models describe plant fitness as dependent on plant interaction intensity and density of surrounding individuals (Hart et al. 2018). Plant interaction intensity between two individuals further depends on the identity of the two individuals, as well as the distance between them (Purves and Law 2002). This framework allows to infer individual productivity based on the identity, density and distance of the surrounding plants.

In this project, we used this theory to predict optimal sowing patterns in intercropping systems. Specifically, we developed a spatially-explicit model of crop productivity based on the identity and abundance of an individual’s neighbours and the distance between them. We used ecological interaction models based on density and distance to quantify the competitive abilities of the considered crops, and how these competition coefficients vary with distance (Vandermeer 1986, Hart et al. 2018). This model allowed us to find the spatial arrangement that maximizes either population-level yield, i.e. crop yield for each intercropped species, or community-level yield, i.e. the combined yield of all the species forming the intercrop. The latter can be expressed relative to the expected yield based on the corresponding monoculture yields using the Land Equivalent Ratio (LER). We focused on three species in this study, namely one cereal (*Avena sativa)*, one legume (*Lupinus angustifolius)*, and one Brassicaceae (*Camelina sativa)*. Cereal-legume intercrops are generally very efficient intercropping combinations due to complementary nitrogen acquisition strategies (Hauggaard-Nielsen et al. 2010). Intercrops with non-legumes are much less studied; here we chose a Brassicaceae, as this family shows several promising aspects for successful intercropping, such as allelopathic activity (Engbersen et al. 2021). By modelling crop yield from simple density and distance experiments, this research offers an easy way to optimize crop combinations and sowing patterns for minimal competitive interactions and maximal crop yield.

## 2. Material and methods

### 2.1. Study site

The density and distance experiments took place in an outdoor experimental garden at the Aprisco de Las Corchuelas research station (39.8133 N, 6.0003 W, 350 m a.s.l.) in Torrejón el Rubio, Cáceres, Spain. The experimental garden consisted of 170 square boxes of 0.25 m^2^ with a depth of 40 cm. Beneath 40 cm the boxes were open, allowing unlimited root growth in the underlying soil. The boxes were embedded into larger beds of 10 × 1 m^2^. Inside a bed, plots were separated from each other by metal frames. Each box was filled down to 30 cm with standard, not enriched, agricultural soil coming from the local region. This soil was composed of 78% sand, 20% silt, and 2% clay, and contained 0.05% nitrogen, 0.5% carbon, and 253 mg total P/kg. The mean pH was 6.30.

The experimental garden was irrigated during the growing season in order to assure survival of the crops during drought periods. Irrigation was automated using soil moisture sensors (PlantCare Ltd., Switzerland); whenever a set dry threshold was reached, irrigation started and added water up to the set target value. During plant germination and establishment, i.e. until two months after sowing, the irrigation was configured for a dry threshold of soil moisture of 40% of field capacity, with a target of 60% of field capacity. After plants were properly established, irrigation was reduced to a dry threshold of 17% of field capacity, with a target of 25% of field capacity.

### 2.2. Crop species

We used three crop species belonging to three different phylogenetic groups with varying functional characteristics: *Lupinus angustifolius* (lupin, legume), *Camelina sativa* (false flax, herb [superrosids]), and *Avena sativa* (oat, C3 grass). We chose these species because they are representative of three large groups of crops currently used in agriculture in Europe, namely cereals, legumes, and oilseed herbs. Cereals and legumes are commonly used in crop mixtures due to complementary use of nitrogen and facilitation in phosphorus uptake (Engbersen et al. 2021). *Avena* is a grass that usually reaches 70 cm in height, with a mean mass per seed of 0.024 grams (Chen et al. 2020). Cereals are furthermore known to be strong competitors for soil nitrogen and are characterized by a high growth rate at early stages (Andrew et al. 2015). *Lupinus* is a legume that grows until an average height of 55 cm, with a mean mass per seed of 0.16 grams (Chen et al. 2020). As a legume, it is able to fix atmospheric N2 and can also facilitate P mobilization by releasing carboxylates into the rhizosphere (Li et al. 2003).

Finally, *Camelina* is a Brassicaceae, which generally grows until 50 cm with an average mass per seed of 0.00092 grams (Chen et al. 2020). Furthermore, Brassicaceae are known to possess allelopathic activity, which can be beneficial to manage pests and diseases or even promote plant growth (Engbersen et al. 2021).

### 2.3. Interaction experiments

Experimental treatments consisted of single plots with either (1) one solitary individual, (2) density experiments with a varying number of neighbors, (3) distance experiments with a varying distance of neighbors (Fig. 1) and (4) test communities.

**Figure 1:**
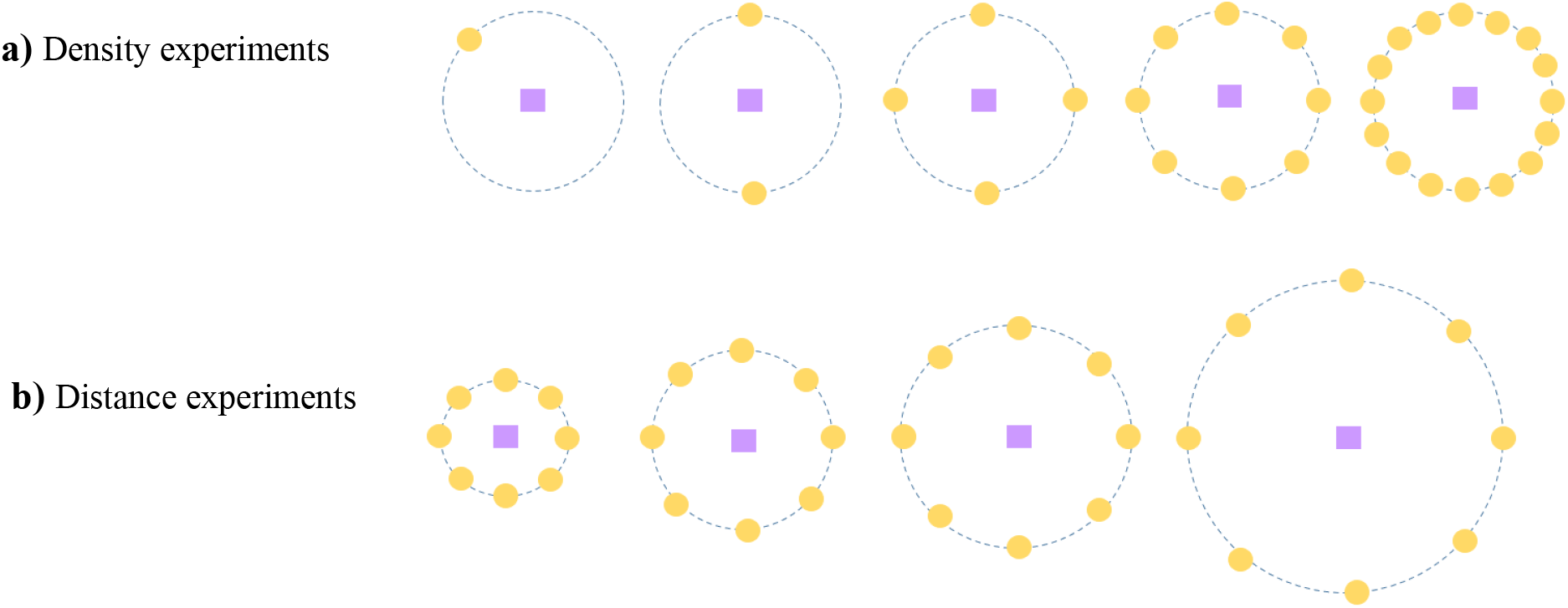
Schematic representations of the density (a) and distance (b) experiments, with a heterospecific pair of species.

#### Density experiments

The density experiments consisted of one focal plant surrounded by none, one, two, four or eight neighbouring individuals, either conspecific (i.e. of the same species) or heterospecific (i.e. of a different species). Neighbours were all planted at the same distance from the focal individual (8 centimeters). We chose this distance as it is the smallest of the standard between-row distances used for cereals (Boström et al. 2012).

#### Distance experiments

Distance experiments consisted of one focal individual surrounded by eight individuals, either conspecific or heterospecific, placed at four different distances from the focal (5, 8, 12 or 20 cm).

#### Test communities

In order to validate the performance of our models, we set up test communities in the above-described plots. These test communities consisted of (i) row alternations of 2 and 3 species with the same plant densities per species and with 8 cm row spacing, (ii) row alternations of 2 and 3 species with varying plant densities per species and with 8 cm row spacing, (iii) homogeneous mixing of 2 and 3 species with 8 cm within- and between-row spacing, and (iv) random arrangements of 3 species with 8 cm within- and between-row spacing. All the possible combinations of two species were planted (i.e. Avena-Lupinus, Avena-Camelina, and Camelina-Lupinus). In the plots with the same plant densities, all species had a density of 150 individuals m^−2^. In the varying densities plots, we kept *Lupinus* with 150 individuals m^−2^, doubled the density of *Avena* (300 individuals m^−2^), and tripled the density of *Camelina* (450 individuals m^−2^), in order to get closer to standard monoculture sowing densities.

We replicated the density, distance and test experiments two times and the single plant plots three times. Plots were fully randomized within and among the beds, with the single plots and the test plots separated from the density and distance plots in order to avoid potential shading of the single individuals. Sowing was conducted by hand on the 2^nd^ and 3^rd^ of February 2019. Germination was evaluated two weeks after sowing and missing individuals were then resown. After that, the number of individuals and focal growth were evaluated every week. *Lupinus* individuals had a particularly low germination rate, due to pest presence. We therefore applied a systemic insecticide (Epik, SIPCAM INAGRA, S.A. [acetamiprid 20%]) three times on all the plots in order to allow germination and growth of the Lupinus individuals.

Until mid-April, dead focal plants were replaced using a pool of extra individuals sown at the beginning of the experiment in a separate bed. After mid-April, considering the size and phenological stage of the plants, replacing and transplanting focal individuals was stopped. Dead neighbouring plants were still replaced using individuals from the extra pool. However, from May onwards, due to the large root system of the plants, transplanting individuals had a very low success rate. Therefore, a neighbouring individual missing for more than half of the experiment was not taken into account for the final number of neighbours.

All plots were manually weeded weekly.

#### Data collection

Harvest was conducted manually in June – i.e. between 124 and 138 days after sowing – when the seeds of the focal species were mature. Aboveground biomass and grain yield of the focal and the neighbours were determined in each plot. Plants were clipped right above the soil surface, the focal individual was separated from the neighbours and seeds were separated from the vegetative parts. In the test communities, plants were separated per species, and seeds were separated from the vegetative parts. Seeds were threshed, sun-dried for five days and then weighed, while vegetative biomass was weighed after drying in an oven at 80°C for 72 hours.

### 2.4. Data analyses

#### 2.4.1. Productivity models and interaction coefficients

To describe the productivity of the focal plant, we used the Beverton-Holt model, which is used to describe annual plant competitive population dynamics in the field (Beverton and Holt 1957, Hart et al. 2018):

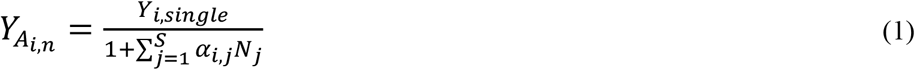

where 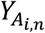 is the productivity (often referred to as fecundity in the plant coexistence framework) (in grams) of the n^th^ focal individual *A_i,n_* of species *i* in the presence of a specified set of neighbours; *Y_i,single_* is the intrinsic productivity (i.e. in the absence of competitors) (in grams) of an individual of species *i; α_i,j_* is the interaction coefficient that represents the direct impact (positive or negative) of species *j* on the focal individual of species *i; N_j_* is the abundance of species *j;* and the summation sign indicates that the sum is taken of all species (S) surrounding the focal individual. If *N*_i_ > 0, *α_i,i_* is included in the sum and therefore represents intraspecific competition.

##### α dependency on distance

Interaction coefficients have been shown to be strongly dependent on interplant distance; that is, as the interplant distance increases, the effect of competition (or facilitation) is expected to decrease. Therefore, the interaction coefficients were calculated as follows (Vandermeer 1984, 1986):

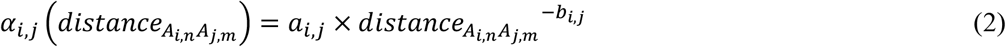

where 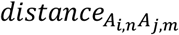 is the distance between the n^th^ individual *A_i,n_* from species *i* and the m^th^ individual *A_j,m_* from species *j; a_i,j_* is a constant representing the intensity of the interaction effect between species *i* and *j*, and *b_i,j_* a constant representing the rate at which that effect decreases with increasing interplant distance.

When combining equations (1) and (2), we obtain the following spatially-explicit plant productivity model (see Fig. 2 for a schematic representation):

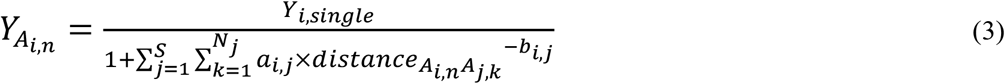

**Figure 2:**
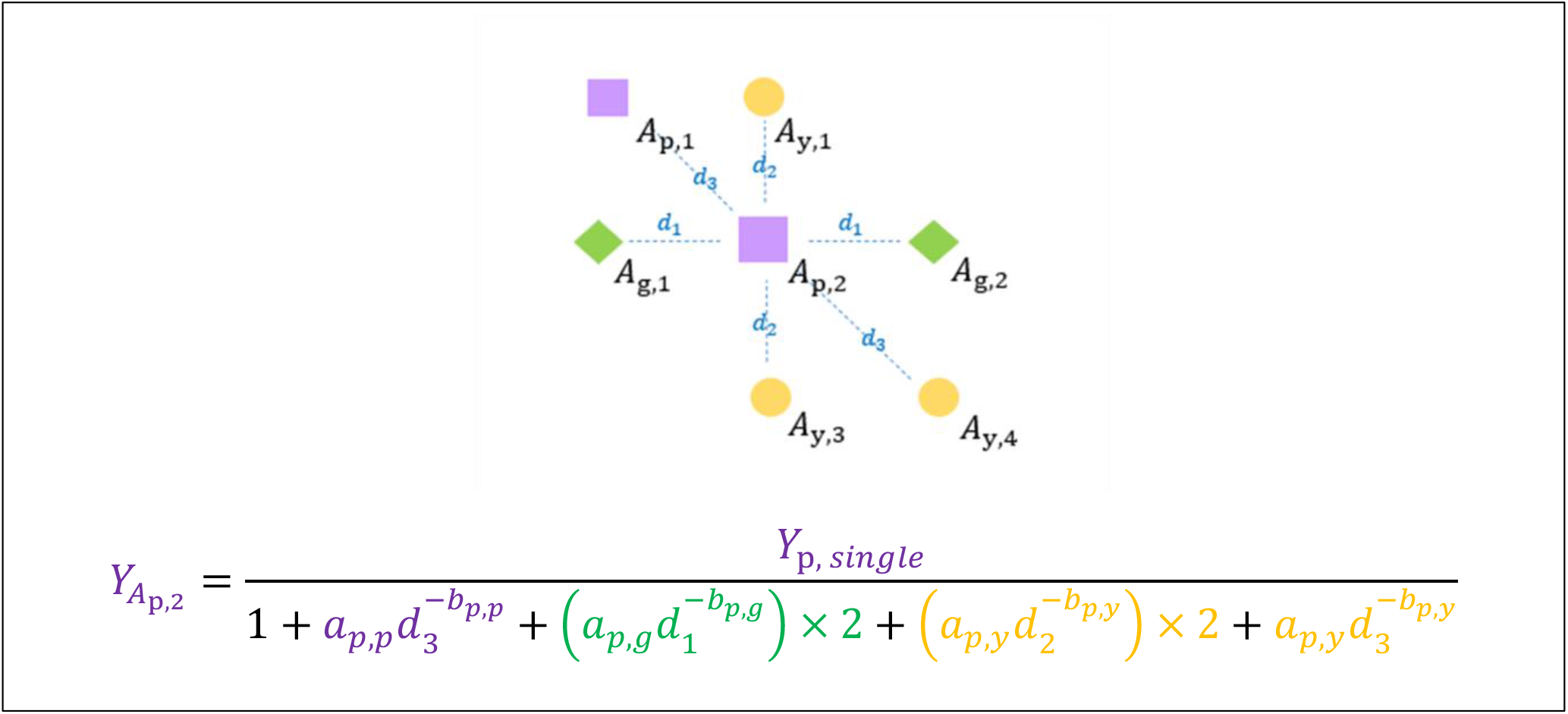
Example of the calculation for the yield of individual *A*_p,2_ using equation (3), with three species (yellow *y*, purple *p*, and green *g*). The letters above the dashed blue lines represent the distances between two individuals.

The productivity of species growing in isolation (*Y_i,single_*) was determined by taking the average yield of the three single individuals per species. Then, we used non-linear least squares (nls) models with the *stats* package in R (R Core Team 2019) to fit the equation (3) to the density and distance data obtained from the field experiment to determine the *a* and *b* coefficients for each conspecific and heterospecific pairwise combination of species. Starting values for the nls models were defined by first fitting a nls Levenberg-Marquardt model (nlsLM, *minpack.lm* package) and reusing the obtained parameters as starting values for the nls model.

##### Model improvement with an intra-specific higher order interaction term

In cases where the fit of the model was poor (R^2^ < 0.50) (Table S1), we tried adding the intra-specific second order interaction term 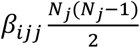 to the denominator (Mayfield and Stouffer 2017) and kept it if this term improved the fit of the model to the data points. This term represents the intraspecific crowding effect which occurs among N individuals of species *j* on the focal individual of species *i*, with the strengths of these effects given by *β_ijj_*.

#### 2.4.2. Species-level yield and LER simulations

We used the previously-described model in a spatially-explicit approach to calculate the yield per species at the field scale. In this study we consider up to three species, referred to in this example as A, B and C. For example, consider a matrix *a* of dimension *n* × *m* of individuals A, B and C (see example in Fig. 3). Each individual is defined by its position (*k,l)* in the matrix, as well as by its species identity value *a*[*k,l*]. In a first step, for each individual on location (*k,l)*, we calculated its distances to every other individual at location *(p,q)*, with *p* and *q* ranging from 1 to *n* and 1 to *m*, respectively, using the Euclidian distance 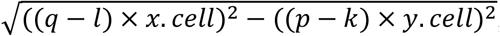, where *x. cell* and *y. cell* represent the distance between the rows and columns of the matrix *a* (Fig. 3). Then, for the individual (*k,l)*, we selected all the neighboring individuals satisfied the condition that their distance to (*k,l)* was smaller than a pre-defined distance (*d.lim*). Among the selected neighbours, we further selected only the eight closest neighbours – the Moore neighborhood (Yitbarek and Vandermeer 2017) – that were directly connected to individual *(k,l)* (i.e. without any other individual in the straight line between them and (*k,l)*). We chose to select these eight closest neighbours as including individuals further away did not improve the fit of our model (see section 2.4.3). Based on these selected neighbours, we calculated the yield of individual (*k,l)* using equation (3). This process was repeated for every individual in the plant matrix *a*. Finally, we summed the yield of each species; to avoid any edge effect at the boundary of the matrix, we only considered individuals that were not affected by the matrix boundaries (i.e. we excluded a buffer zone of potential edge effects ranging from rows 1 + *buffer* and *n* − *buffer* to columns 1 + *buffer* and *m* − *buffer*, see Fig. SX). We then divided the obtained yield by the “area” (*n* − 2 × *buffer*) × *y. cell* × (*m* − 2 × *buffer*) × *x. cell* of the buffered plant matrix *a* to get a measure of yield in g.m^−2^.

**Figure 3:**
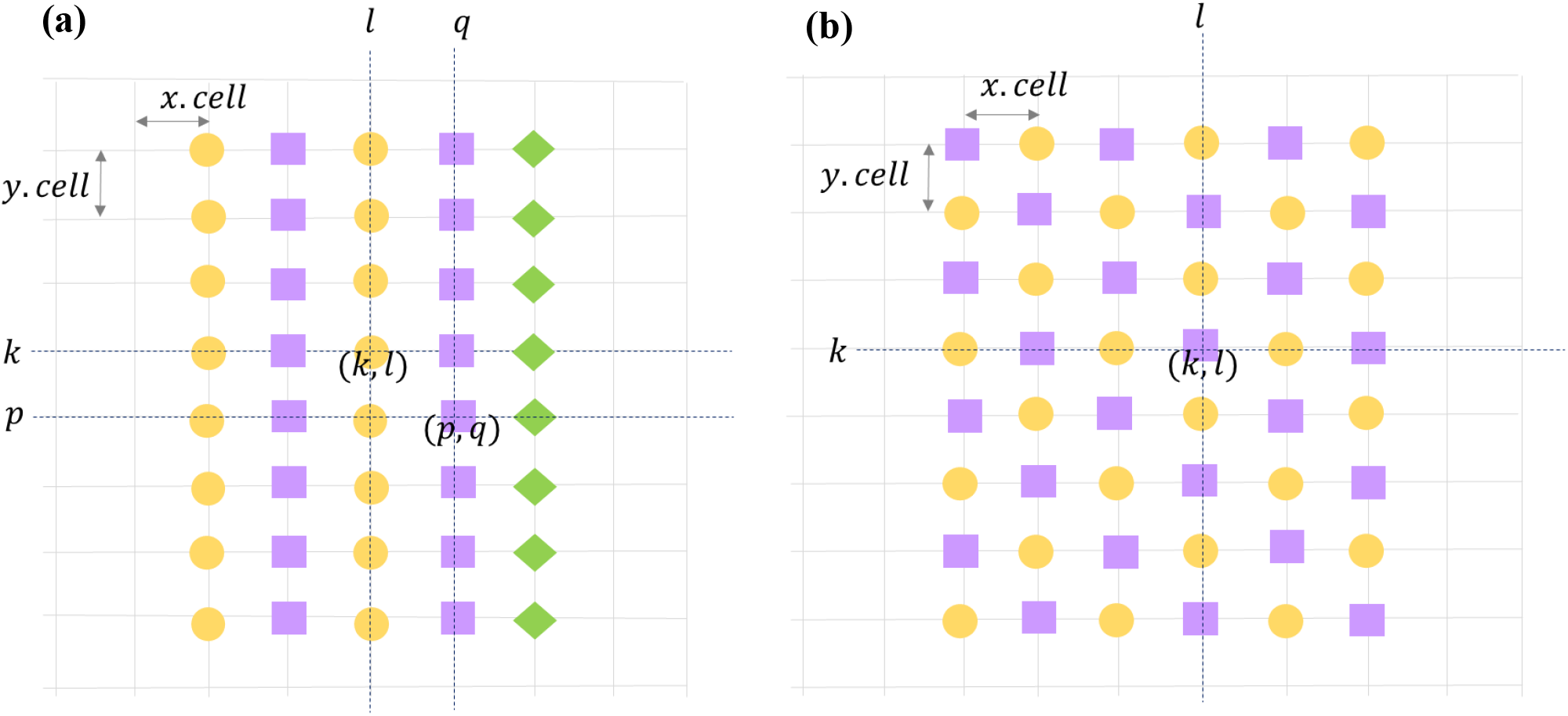
Example of plant matrices showing a row configuration with three species (a) and a homogeneous mixture with two species (b). *(k,l)* designate the coordinates of an individual in the matrix; *x.cell* and *y.cell* represent the inter-column and inter-row distance.

##### LER

The community-level yield was assessed by calculating the Land Equivalent Ratio (LER):

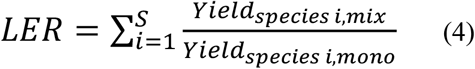

where *Yield_species i,mix_* represents the yield (g.m^−2^) of species *i* in mixtures, and *Yield_species i,mono_* the yield (g.m^−2^) of species *i* in monocultures. The ratio *Yield_species i,mix_/Yield_species i,mono_* for a given species *i* represents its partial LER. We considered two options for *Yield_species i,mono_*. Firstly, we simulated the monoculture yield with the spacing that we used for our test communities (i.e. inter-row of 8 cm, intra-row of 5 cm). This constituted the standard monoculture yield. Secondly, we selected the highest monoculture yield among intra- and inter-row distances which were varied from 5 to 20 cm in our model. This constituted the optimal calculated monoculture yield.

#### 2.4.3. Model validation

We used our model to simulate the yield per species of the test communities from our field experiment, and subsequently compared the simulated values to the actual measured values using linear models for each species, with plot row as random factor. We also used these comparisons to test different values of *d.lim* and subsequently selected the value of *d.lim* for which the linear model coefficients were significant and closest to one.

#### 2.4.4. Considered configurations

We considered four types of spatial plant configurations for our simulations (see Fig. 4): row alternation with equal densities of species, homogeneous mixing, row alternation with different densities of species and random placement. We simulated every pair of two-species mixtures and the three-species mixture for all the configurations above, except for the random placement where we only considered the three-species mixture. For the row alternation with three species, we added an extra combination in the following pattern: Avena-Lupinus-Camelina-Lupinus. Finally, the density variations were followed as indicated in Table 1. We simulated species-level yield and LER (standard and optimal) for each of these configurations with inter- and intra-row distance varying from 5 cm to 20 cm, i.e. the distances covered by our experiments.

**Table 1:** Table showing the density ratios of the considered configurations. For instance, in the density 2:1, there is twice as many Camelinas as there are Avenas. *NA:* not applicable.

| *Species combinations* | *Density 2:1* |  |  | *Density 3:2:1* |  |  |
| --- | --- | --- | --- | --- | --- | --- |
|  | Camelina | Avena | Lupinus | Camelina | Avena | Lupinus |
| Avena-Camelina | 2 | 1 | NA | NA | NA | NA |
| Avena-Lupinus | NA | 2 | 1 | NA | NA | NA |
| Camelina-Lupinus | 2 | NA | 1 | NA | NA | NA |
| Avena-Camelina-Lupinus | 2 | 1 | 1 | 3 | 2 | 1 |
| Avena-Lupinus-Camelina-Lupinus | 2 | 1 | 1 | 3 | 2 | 1 |

**Figure 4:**
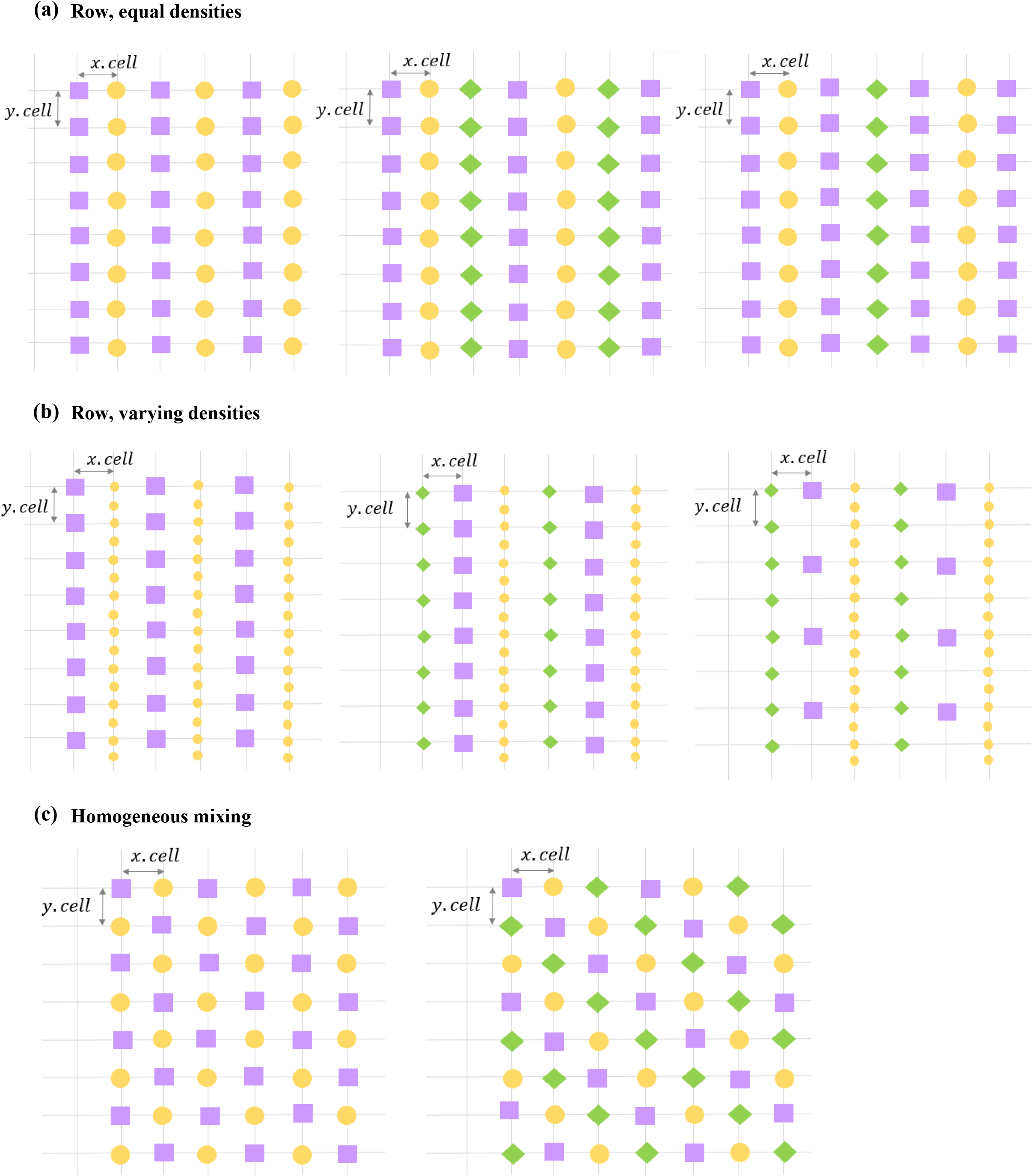
Schematic representations of the spatial configurations considered for the simulations.

## 3. Results

### 3.1 Growth model coefficients

The coefficient of determination, R^2^, for the growth models estimating individual reproductive productivity as a function of neighborhood identity and distance was generally high, i.e. higher than 0.70 for seven out of nine combinations (Fig. 5, Table S1). Only the fit of Camelina with Camelina was less good (R^2^ = 0.51), while the fit of Camelina with Lupinus was relatively poor (R^2^ = 0.34).

**Figure 5:**
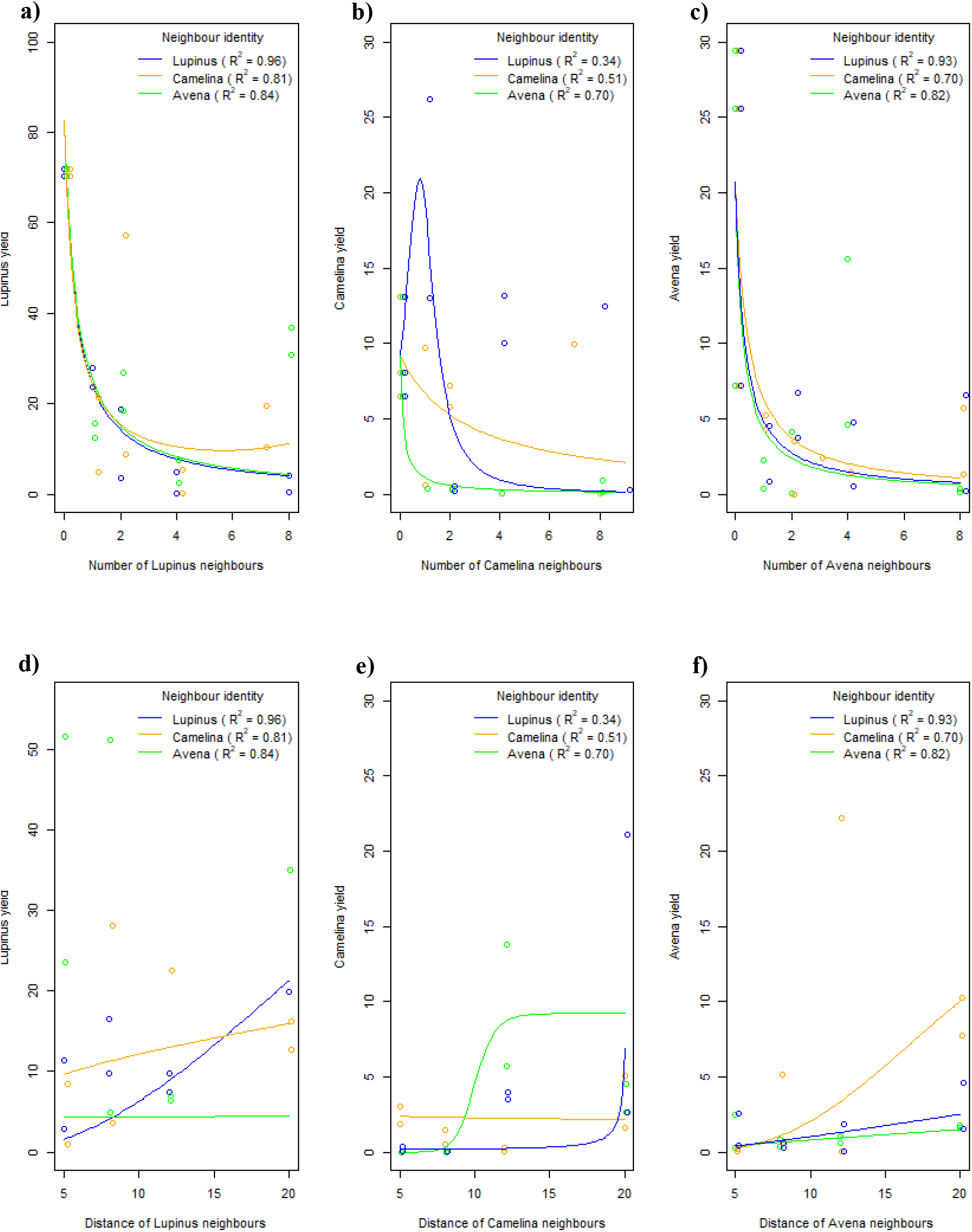
Yield-density (a,b,c) and yield-distance (d,e,f) curves obtained with the equations from Table S2, showing the yield of Lupinus (a,d), Camelina (b,e), and Avena (c,f). See Table S2 for the model equations.

### 3.2 Model validation with test communities

The estimated yield for the test communities based on the individual growth models were significantly correlated with the observed yield in the case of Avena (linear regression coefficient: 0.262, p-value = 0.0033) and Lupinus (linear regression coefficient: 0.255, p-value = 0.045), but only slightly significant in the case of Camelina (linear regression coefficient: 0.802, p-value = 0.0785).

### 3.3 Simulations and optimizations

#### Optimized monocultures

For each simulated monoculture, we determined the optimal inter- and intra-row distance for maximal yield at the field scale (Table 2, Fig S1). Avena and Camelina reached the maximum population yield at the smallest simulated distance (5 cm), i.e. the highest densities encompassed in the scope of this study (400 individuals.m^−2^). This increase in density led to an increase in population yield of 27% for Avena and 131% for Camelina compared to the standard density (8 cm). The optimal spatial parameters for Lupinus were inter- and intra-row distances of 6 and 6 cm, respectively, which corresponds to a density of 278 individuals.m^−2^. This allows for an increase in yield of 4.7% compared to the standard density.

**Table 2:** Usual and optimal inter-row and intra-row distances (in cm), and corresponding population level yield (in g.m^−2^) for each species monocultures.

| Species monoculture | Standard inter-row distance (cm) | Standard intra-row distance (cm) | Standard yield (g.m<sup>-2</sup>) | Optimal inter-row distance (cm) | Optimal intra-row distance (cm) | Optimal yield (g.m<sup>-2</sup>) |
| --- | --- | --- | --- | --- | --- | --- |
| Avena | 8 | 5 | 153 | 5 | 5 | 195 |
| Camelina | 8 | 5 | 184 | 5 | 5 | 425 |
| Lupinus | 8 | 5 | 813 | 6 | 6 | 851 |

#### Optimized mixtures

Using the model simulations, we determined spatial configurations (i.e. homogeneous vs. row pattern) and sowing densities (optimal inter- and intra-row distances) that maximized community-level yield for each mixture (Table 3, Fig S2, S3, S4). Increases in total yield in these optimized configurations compared to the ones we used in the test communities ranged from 7% for the 3-species random placement to 219% for the Avena-Camelina row pattern.

**Table 3:** Optimal inter-row and intra-row distances (in cm) for each species combinations and spatial configurations.

| Species combination | Type of configurations | Density | Optimal inter-row distance (cm) | Optimal intra-row distance (cm) | Standard total yield (g.m<sup>-2</sup>) | Optimal total yield (g.m<sup>-2</sup>) | Yield increase (%) |
| --- | --- | --- | --- | --- | --- | --- | --- |
| Avena-Camelina | Row | Equal | 11 | 5 | 172 | 548 | 219 |
| Avena-Lupinus | Row | Equal | 5 | 5 | 467 | 709 | 52 |
| Camelina-Lupinus | Row | Equal | 5 | 6 | 538 | 841 | 56 |
| Avena-Camelina | Homogeneous | Equal | 10 | 10 | 98 | 158 | 61 |
| Avena-Lupinus | Homogeneous | Equal | 5 | 5 | 506 | 832 | 64 |
| Camelina-Lupinus | Homogeneous | Equal | 5 | 5 | 579 | 938 | 62 |
| Avena-Camelina | Row | 2:1 | 11 | 5 | 247 | 545 | 121 |
| Avena-Lupinus | Row | 2:1 | 5 | 5 | 395 | 600 | 52 |
| Camelina-Lupinus | Row | 2:1 | 5 | 5 | 742 | 1044 | 41 |
| Avena-Camelina-Lupinus | Row | Equal | 12 | 5 | 385 | 506 | 31 |
| Avena-Camelina-Lupinus | Homogeneous | Equal | 5 | 5 | 361 | 625 | 73 |
| Avena-Camelina-Lupinus | Row | 2:1 | 10 | 5 | 451 | 608 | 35 |
| Avena-Camelina-Lupinus | Row | 3:2:1 | 10 | 5 | 372 | 550 | 48 |
| Avena-Camelina-Lupinus | Random | Random | 5 | 8 | 442 | 474 | 7 |
| Avena-Lupinus-Camelina-Lupinus | Row | Equal | 5 | 5 | 462 | 708 | 53 |
| Avena-Lupinus-Camelina-Lupinus | Row | 2:1 | 5 | 5 | 526 | 763 | 45 |
| Avena-Lupinus-Camelina-Lupinus | Row | 3:2:1 | 5 | 5 | 533 | 696 | 31 |

In all the row configurations, the optimal intra-row distance was generally small (almost always 5 cm), while the optimal inter-row distance showed more variability. In particular, the optimal inter-row distance between Avena and Camelina was larger than between Avena and Lupinus or Camelina and Lupinus: it was either 10 (in homogeneous mixing), 11 (in 2-species row pattern) or 12 cm (in 3-species row pattern). When looking at the homogeneous mixing, the optimal inter- and intra-row distances were the smallest possible in the case of Avena-Lupinus (5 cm), Camelina-Lupinus (5 cm), and the three species mixtures (5 cm) but larger for the Avena-Camelina mixture (10 cm).

#### LER gain

Next, we compared the yield of mixtures to those of monocultures using LER. Increases in LER when the mixtures were spatially optimized compared to the distances used in the test communities ranged from 8% for the 3-species random placement to 142% for the Camelina-Lupinus row pattern (Table 4).

**Table 4:**
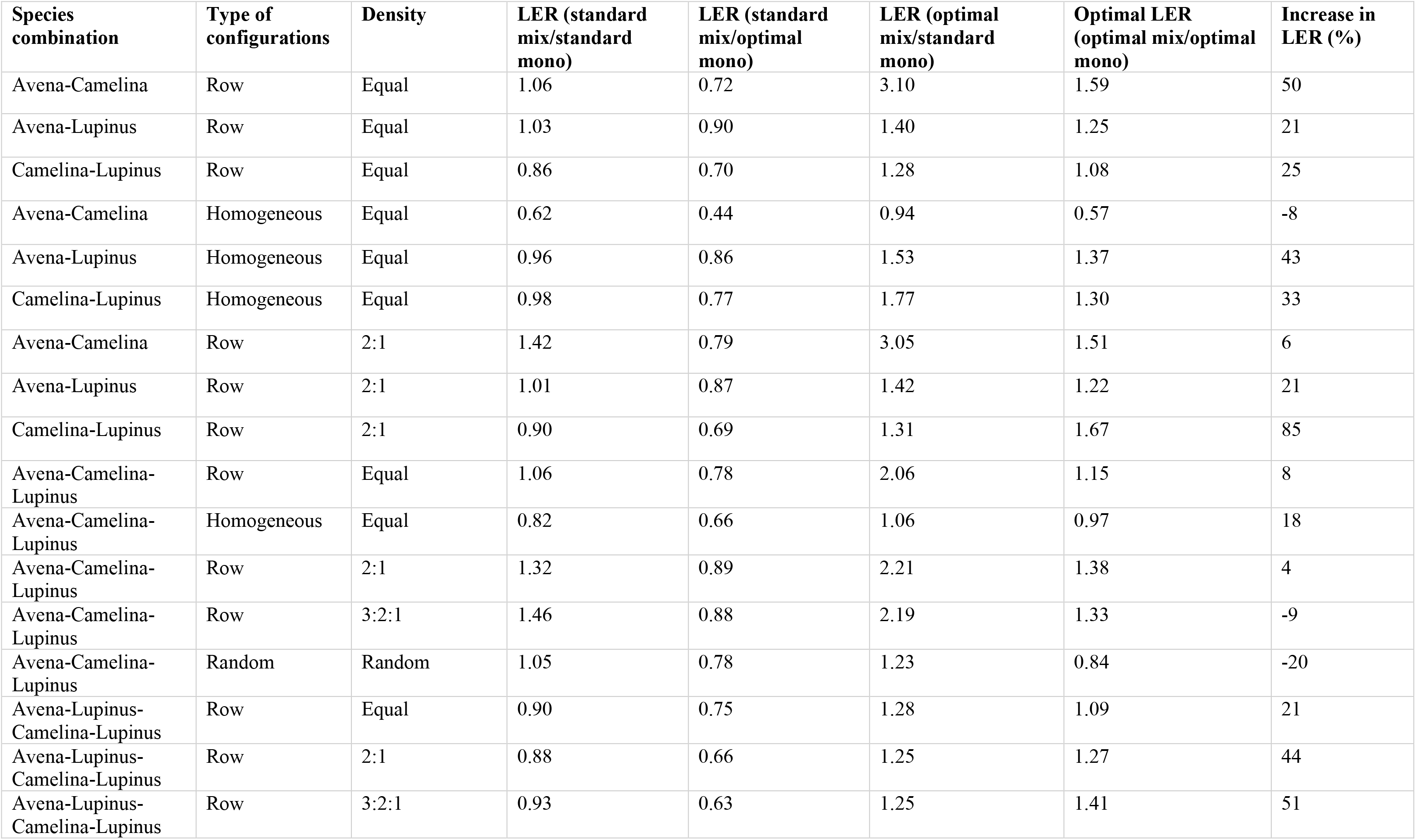
Standard and optimal LERs for each species combination and spatial configuration. Standard monoculture and mixture refer to the yield simulated with the spacing that we used for our test communities; optimal monoculture and mixture refer to the highest selected yield among intra- and inter-row distances varying from 5 to 20 cm.

| Species combination | Type of configurations | Density | LER (standard mix/standard mono) | LER (standard mix/optimal mono) | LER (optimal mix/standard mono) | Optimal LER (optimal mix/optimal mono) | Increase in LER (%) |
| --- | --- | --- | --- | --- | --- | --- | --- |
| Avena-Camelina | Row | Equal | 1.06 | 0.72 | 3.10 | 1.59 | 50 |
| Avena-Lupinus | Row | Equal | 1.03 | 0.90 | 1.40 | 1.25 | 21 |
| Camelina-Lupinus | Row | Equal | 0.86 | 0.70 | 1.28 | 1.08 | 25 |
| Avena-Camelina | Homogeneous | Equal | 0.62 | 0.44 | 0.94 | 0.57 | -8 |
| Avena-Lupinus | Homogeneous | Equal | 0.96 | 0.86 | 1.53 | 1.37 | 43 |
| Camelina-Lupinus | Homogeneous | Equal | 0.98 | 0.77 | 1.77 | 1.30 | 33 |
| Avena-Camelina | Row | 2:1 | 1.42 | 0.79 | 3.05 | 1.51 | 6 |
| Avena-Lupinus | Row | 2:1 | 1.01 | 0.87 | 1.42 | 1.22 | 21 |
| Camelina-Lupinus | Row | 2:1 | 0.90 | 0.69 | 1.31 | 1.67 | 85 |
| Avena-Camelina-Lupinus | Row | Equal | 1.06 | 0.78 | 2.06 | 1.15 | 8 |
| Avena-Camelina-Lupinus | Homogeneous | Equal | 0.82 | 0.66 | 1.06 | 0.97 | 18 |
| Avena-Camelina-Lupinus | Row | 2:1 | 1.32 | 0.89 | 2.21 | 1.38 | 4 |
| Avena-Camelina-Lupinus | Row | 3:2:1 | 1.46 | 0.88 | 2.19 | 1.33 | -9 |
| Avena-Camelina-Lupinus | Random | Random | 1.05 | 0.78 | 1.23 | 0.84 | -20 |
| Avena-Lupinus-Camelina-Lupinus | Row | Equal | 0.90 | 0.75 | 1.28 | 1.09 | 21 |
| Avena-Lupinus-Camelina-Lupinus | Row | 2:1 | 0.88 | 0.66 | 1.25 | 1.27 | 44 |
| Avena-Lupinus-Camelina-Lupinus | Row | 3:2:1 | 0.93 | 0.63 | 1.25 | 1.41 | 51 |

We show that for almost all species combinations (except Avena-Lupinus), the maximum LER of the row pattern is higher than the maximum LER of the corresponding homogeneous mixture (Fig. 7). This means that mixtures are more advantageous for productivity when crops are arranged in a row alternation than when they are completely mixed. For instance, the alternation of rows for Avena-Camelina led to a 2.8 times higher LER than the homogeneous pattern. Similarly, for the three-species mixture, the LER for row alternation was 1.2 times higher than for the homogeneous mixing. The only exception is the Avena-Lupinus mixture, where the homogeneous mixture led to a slightly higher LER (+10%) than the row alternation with the same density of species.

Moreover, we found that increasing plant density did not necessarily lead to more efficient crop mixtures (Fig. 7). For instance, in the case of Camelina-Lupinus, reducing sowing density of Lupinus from 200 to 100 seeds.m^−2^ led to a 55% increase in LER. This was also the case for the three-species mixture Avena-Camelina-Lupinus, where decreasing the densities of both Lupinus and Avena from 66 to 33 seeds.m^−2^ increased LER by 20%. Similarly, for the Avena-Lupinus-Camelina-Lupinus mixture, reducing the density of Lupinus from 200 to 100 seeds.m^−2^ and the density of Avena from 100 to 50 seeds.m^−2^ led to a 16% LER increase. Further reducing the density of Lupinus to 66 seeds.m^−2^ increased LER by 11%.

Finally, 3-species mixtures did not show higher LER than 2-species mixtures; specifically, Avena-Camelina and Camelina-Lupinus were able to reach higher LERs than the three species together (+13% for Avena-Camelina and +18% for Camelina-Lupinus). The mixture of Avena-Lupinus reached a similar LER value than the three-species mixture.

When decomposing LER into partial LERs (Fig. 7), in most cases the increase in total LER was substantially driven by an increase in the contribution of Camelina. This is notably true for the Avena-Camelina mixture, where the partial contribution of Camelina increased by 420% in the optimized row pattern (i.e. 11 cm inter-row) compared to the homogeneous mixing. Similarly, in the three-species mixtures, the contribution of Camelina went from almost 0 in the homogeneous mixing to 0.73 in the optimized row pattern. On the contrary, the partial contribution of Avena to total LER remained mostly similar across mixture combinations and spatial configurations. Finally, the relative contribution of Lupinus was generally high in the Camelina-Lupinus mixture, as well as in the three species homogeneous mixture and the Avena-Lupinus-Camelina-Lupinus with equal densities. Then it drastically decreased by up to 88% in the other three-species configurations, in favor of the increase of the partial contribution of Camelina.

## 4. Discussion

This study demonstrates that models from plant coexistence theory can be used to optimize crop species combinations and spatial configurations in intercropped systems. While the density experiments gave us some clues on which species would perform well together, the distance experiments allowed us to understand how the changes in yield associated with the presence of neighbours varied with distance. Based on these elements, we were able to find the sets of parameters (identity of neighbours, sowing density, distances between individuals) that would optimise intercrop yield (measured as LER) in terms of species combinations and spatial configurations.

### Promising crop combinations

Our results from the density and distance experiments provided helpful clues as to which species benefit from growing together. In particular, we noticed that Avena and Camelina strongly compete with each other: the yield-density curve of Camelina with Avena indeed decreased rapidly and steeply (Fig. 5b). However, the distance curve (Fig. 5e) suggests that there may be a limit after which the competitive effect of Avena on Camelina is weaker. This indicates that Avena and Camelina individuals must be far from each other (i.e. separated by more than 11 cm in our case) for a successful mixture (Fig. S1a-b-c). Few previous studies have investigated the case of Avena-Camelina intercropping; in one study of Camelina intercropped with wheat and with lupin, it was shown that Camelina yields were lower in the wheat mixture than in the lupin mixture, which is consistent with our results and further suggests that cereals are strong competitors of Camelina (Paulsen 2007).

On the contrary, promising crop combinations include Camelina-Lupinus and Avena-Lupinus. First, the yield-density curve of Camelina with Lupinus neighbours suggests that there may be some facilitative effects at a low density of Lupinus (Fig. 5b). This has the potential to be highly beneficial for the species-level yield of Camelina in the case of a lower density of Lupinus, which we can see in our Camelina-Lupinus LER simulations (Fig. 7). Furthermore, the optimal LERs were quite high (Fig. 7); in his study, Paulsen (2007) similarly showed that intercropping of Lupinus and Camelina increased the total relative yield, which suggests that this is a successful, beneficial mixture.

In the case of the Avena-Lupinus combination, the two yield-density curves look very similar (Fig. 5a–c), which could indicate that competition between the species is symmetrical (Weiner et al. 2001). Furthermore, the yield-distance curve of Lupinus surrounded by Avena (Fig. 5d) suggests that Lupinus and Avena could perform well in close proximity. Indeed, our simulation results show that LER for Avena-Lupinus is highest when the two species are as close as possible (Fig. S1d-e-f). It is also the only mixture where the homogeneous mixing performs as well as the row configuration (Fig. 7). This suggests that Avena is not a strong competitor of Lupinus and that there may be a high level of niche differentiation between the two species, which makes Avena-Lupinus a promising crop combination. Intercropping Lupinus with cereals has indeed been demonstrated as a well-performing mixture in several studies, with LERs generally superior than one due to a complementary use of resources, particularly nitrogen (Engbersen et al. 2021).

Surprisingly, contrary to the diversity-productivity relationship (Loreau and Hector 2001), the three-species mixtures did not perform more efficiently than the best two-species mixtures (Fig. 7). Reasons for this lack of increase might be that going from two species to three species is not a substantial enough increase in crop diversity. It may also be that the three species are too similar in terms of growth strategies, or that our model fails to predict plant growth when more than two species are considered (see below).

### Optimal spatial configurations

Our simulations show that, except Avena-Lupinus, all combinations perform better in a row pattern than in a homogeneous mixing (Fig. 7); this may be because individuals are directly surrounded by more conspecific neighbours in the homogeneous mixing than in the row pattern. Since plants generally compete more strongly with individuals from their own species than with individuals from a different species (Adler et al. 2018), they tend to benefit from being surrounded by more heterospecific neighbours than conspecific neighbours. This is consistent with a recent meta-analysis showing that net biodiversity effects (i.e. the increase in yield in mixtures compared to monocultures) were significantly greater in alternate-row than in homogeneously mixed intercrops (Li et al. 2020), and with a recent study demonstrating that alternating rows were more beneficial to increase soybean production while also controlling weed invasions (Cheriere et al. 2020).

Furthermore, our results demonstrate that increasing plant density does not necessarily increase yield (Fig. 7); there is a trade-off between density and productivity, i.e. between a few highly productive plants and many less productive individuals, also known as the constant yield law (Weiner et al. 2001, Weiner and Freckleton 2010). In our case, we show for instance that decreasing the density of Lupinus by a factor 2 or 3 can be beneficial for LER (up to + 29%). This is in agreement with previous research on intercropping with legumes, where legumes usually have a much lower density than the other intercropped species (Bulson et al. 2020). On the contrary, in the two-species mixtures, decreasing the density of Avena did not translate into an increase in LER. This is consistent with most of the cereal intercropping research, which shows that increasing the density of the cereal in mixtures generally leads to higher LERs (Yu et al. 2016, Bulson et al. 2020). Cereals are indeed commonly considered to be strong competitors, notably due to their high growth rate at early life stages and high uptake of nitrogen (Andrew et al. 2015). Furthermore, among the three species we considered in this study, cereals have the longest history of breeding for higher population yield, which is achieved by decreasing, not increasing, individual fitness (Denison 2012, Weiner 2017). Thus, we believe that cereals are adapted to reach high population-level yield, which can explain why they can withstand high densities without hindering their performance.

### Model performance and limitations

When comparing the simulated monoculture yields with actual average yields from the local region, we found similar values for Avena, with yields ranging from 1100 to 1600 kg.ha^−1^ (Sánchez-Martín et al. 2014, “Bienes asegurables 2020”), in line with our simulations (1530 kg.ha^−1^, Table 2). Local yields for Lupinus are generally smaller than what we simulated (8130 kg. ha^−1^) and only reach up to 3000 kg.ha^−1^ in the field (López-Bellido et al. 1994, Lema et al. 2005). However, the studies mentioned mostly consider *Lupinus albus*. Furthermore, our simulations are consistent with results from a crop diversity experiment performed in 2018 with the same crop varieties and experimental conditions, where the yield of Lupinus and Camelina in monocultures had an average of 8840 kg. ha^−1^ and 1160 kg. ha^−1^, respectively (Stefan et al. 2021).

Our simulated values correlated reasonably well with the experimental values for Lupinus and Avena, but only slightly for Camelina (Fig. 6). We believe this is because our model fails to completely capture the behavior of Camelina when surrounded by Lupinus. Indeed, the R-square for the growth model was particularly low (0.34, Table S1), which could be due to the potential facilitation occurring at low density of Lupinus, particularly in the case of one and four Lupinus neighbours (Fig 4). More complex models including higher-order interactions might improve model fits (Martyn et al. 2020). In this study, our model only considered first-order interactions (representing the direct effect of neighbours) and intraspecific second-order interactions (representing the indirect effect of intraspecific crowding) when it improved the fit of the model. Interspecific higher-order interactions, which capture the indirect effects of all heterospecific pairs of species, were not added to the presented simulations in this study. Including interspecific higher order interactions would require more experiments, notably with combinations of three species (Mayfield and Stouffer 2017, Levine et al. 2017). Thus, more experiments and replications are needed to confirm the observed interactions and to improve the models. In particular, we advocate for a higher number of replicates and an expansion of the number of density-distance combinations (i.e. repeating the distance experiments for each density treatment). Finally, interaction coefficients are strongly dependent on local abiotic conditions, such as climate or soil parameters (Hart and Marshall 2013). Thus, the value of the interaction coefficient, but also how it changes with distance, is expected to vary under different abiotic conditions (Holzapfel et al. 2006). Therefore, we emphasize the importance of studying plant–plant interactions over a range of conditions (Callaway 1998). This highlights the need to test and validate our model under different environmental conditions to determine how the competition coefficients might be affected, and whether this changes the conclusions for optimal spatial configurations. Nonetheless, this approach, based on simple and reproducible density and distance experiments, represents a promising way to optimize crop mixtures in terms of both species combinations and spatial configurations.

**Figure 6:**
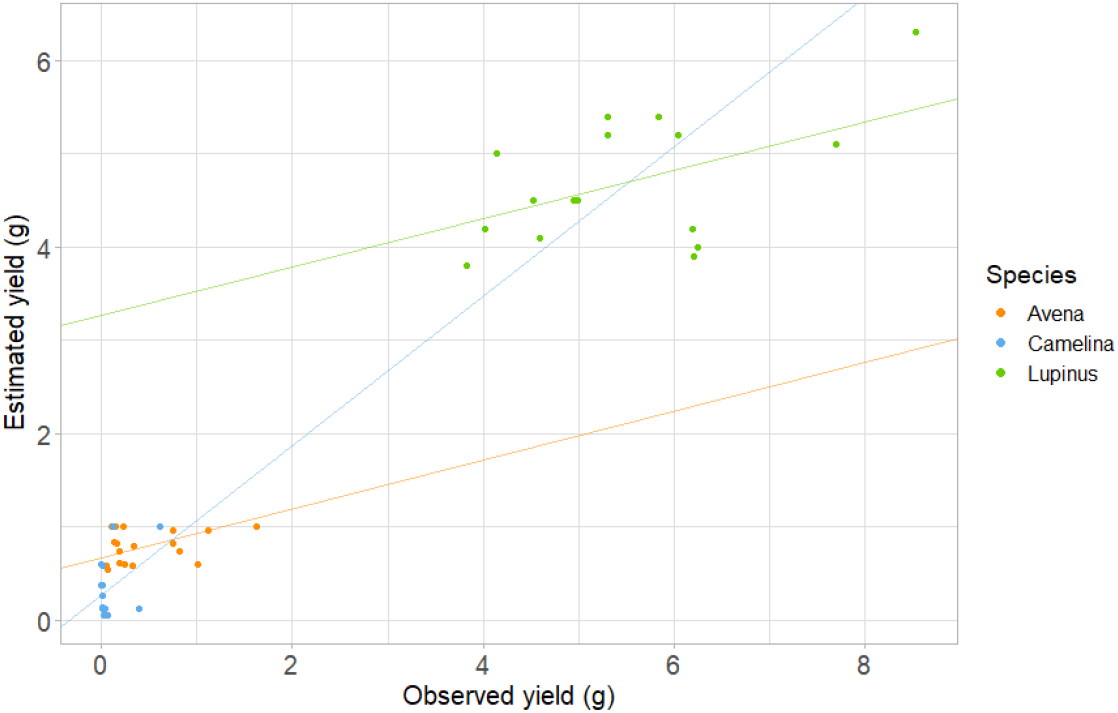
Comparisons between the estimated and observed yields per individuals in the test communities (in g).

**Figure 7:**
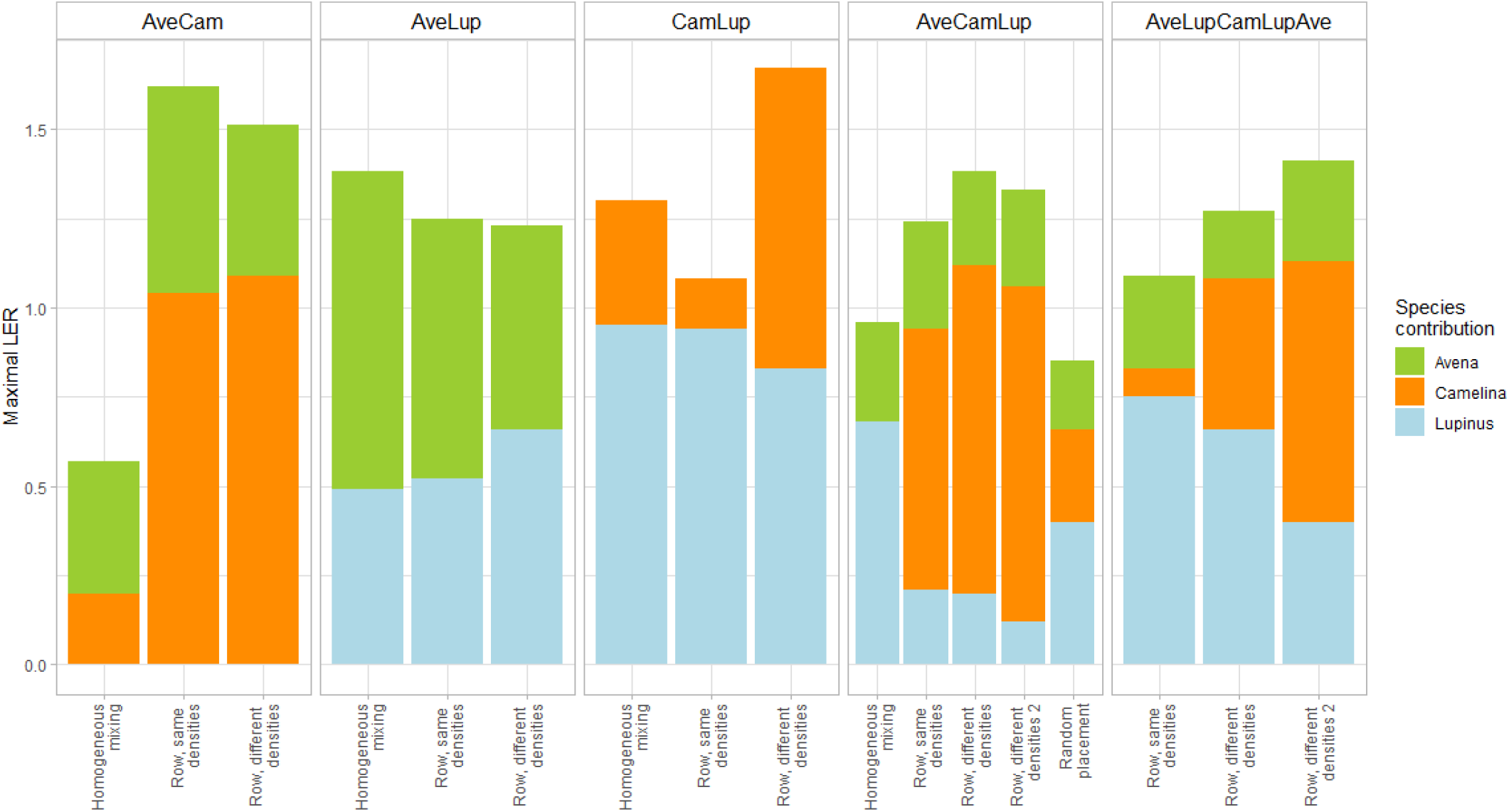
Maximum Land Equivalent Ratios observed for each species combination and spatial configuration. The densities considered in the graph refer to the ones shown in Table 1. AveCam: Avena-Camelina; AveLup: Avena-Lupinus; CamLup: Camelina-Lupinus; AveCamLup: Avena-Camelina-Lupinus; AveLupCamLupAve: Avena-Lupinus-Camelina-Lupinus-Avena.

## Acknowledgements

We would like to thank Sandra Gonzalez Sanchez and Nicola Haggenmacher for their help in the field, and Jacob Usinowicz and Marijn Van de Broek for their help with the modelling part. We also thank the Aprisco de Las Corchuelas Field Station for the use of their facilities. The study was funded by the Swiss National Science Foundation (PP00P3_170645).

## Notes

### Competing Interest Statement

The authors have declared no competing interest.

