## Supplementary Information for "Using spatially-explicit plant competition models to optimise crop productivity in intercropped systems"

Supplementary Information Spatial Modelling

| *Focal* | *Neighbour* | *Equation* | $R^2$ |
|---|---|---|---|
| Lupinus | Lupinus | $Y_{lup} = \frac{82.6}{1 + 178.03 \times distance^{-2.07} N_{lup}}$ | 0.96 |
| Lupinus | Camelina | $Y_{lup} = \frac{82.6}{1 + 3.15 \times distance^{-0.13} N_{cam} - 0.46 N_{cam} \frac{N_{cam} - 1}{2}}$ | 0.81 |
| Lupinus | Avena | $Y_{lup} = \frac{82.6}{1 + 2.3 \times distance^{-0.014} N_{ave}}$ | 0.84 |
| Camelina | Camelina | $Y_{cam} = \frac{9.22}{1 + 0.31 \times distance^{0.09} N_{cam}}$ | 0.51 |
| Camelina | Avena | $Y_{cam} = \frac{9.22}{1 + 2.6 \times 10^{14} \times distance^{-15} N_{ave}}$ | 0.70 |
| Camelina | Lupinus | $Y_{cam} = \frac{9.22}{1 - 0.0019 \times distance^{2.7} N_{lup} + 1.78 N_{lup} \frac{N_{lup} - 1}{2}}$ | 0.34 |
| Avena | Avena | $Y_{ave} = \frac{20.7}{1 + 28.3 \times distance^{-0.96} N_{ave}}$ | 0.82 |
| Avena | Lupinus | $Y_{ave} = \frac{20.7}{1 + 59.9 \times distance^{-1.4} N_{lup}}$ | 0.93 |
| Avena | Camelina | $Y_{ave} = \frac{20.7}{1 + 1438 \times distance^{-3.1} N_{cam}}$ | 0.70 |

**Table S1:** Equations describing the yield of the focal in function of the neighbours density and distance.

Monocultures

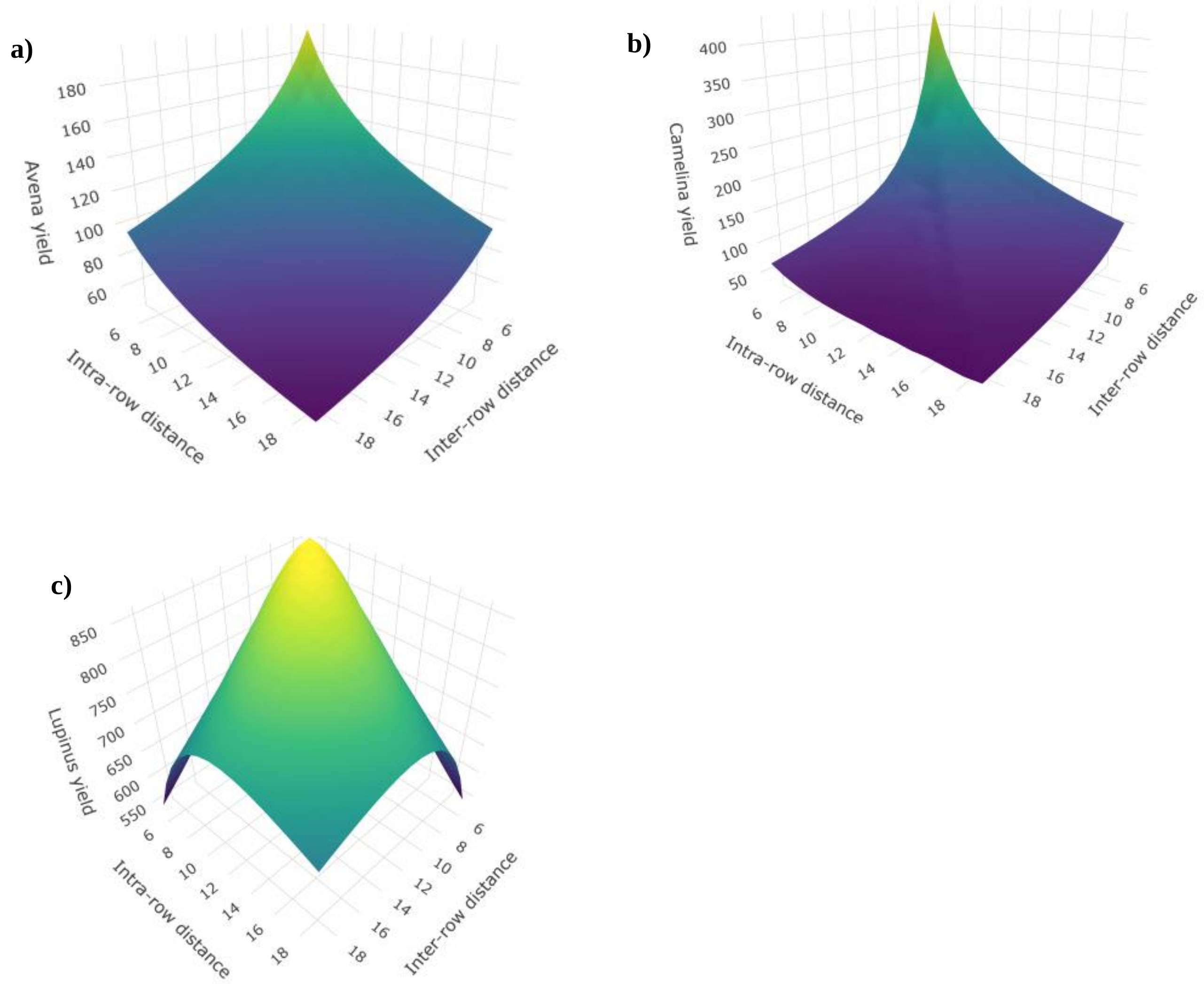


**Figure S1:** Surface plots showing total yield (in g $m^{-2}$) in function of intra- and inter-row distance for monoculture of Avena (a), Camelina (b), and Lupinus (c).

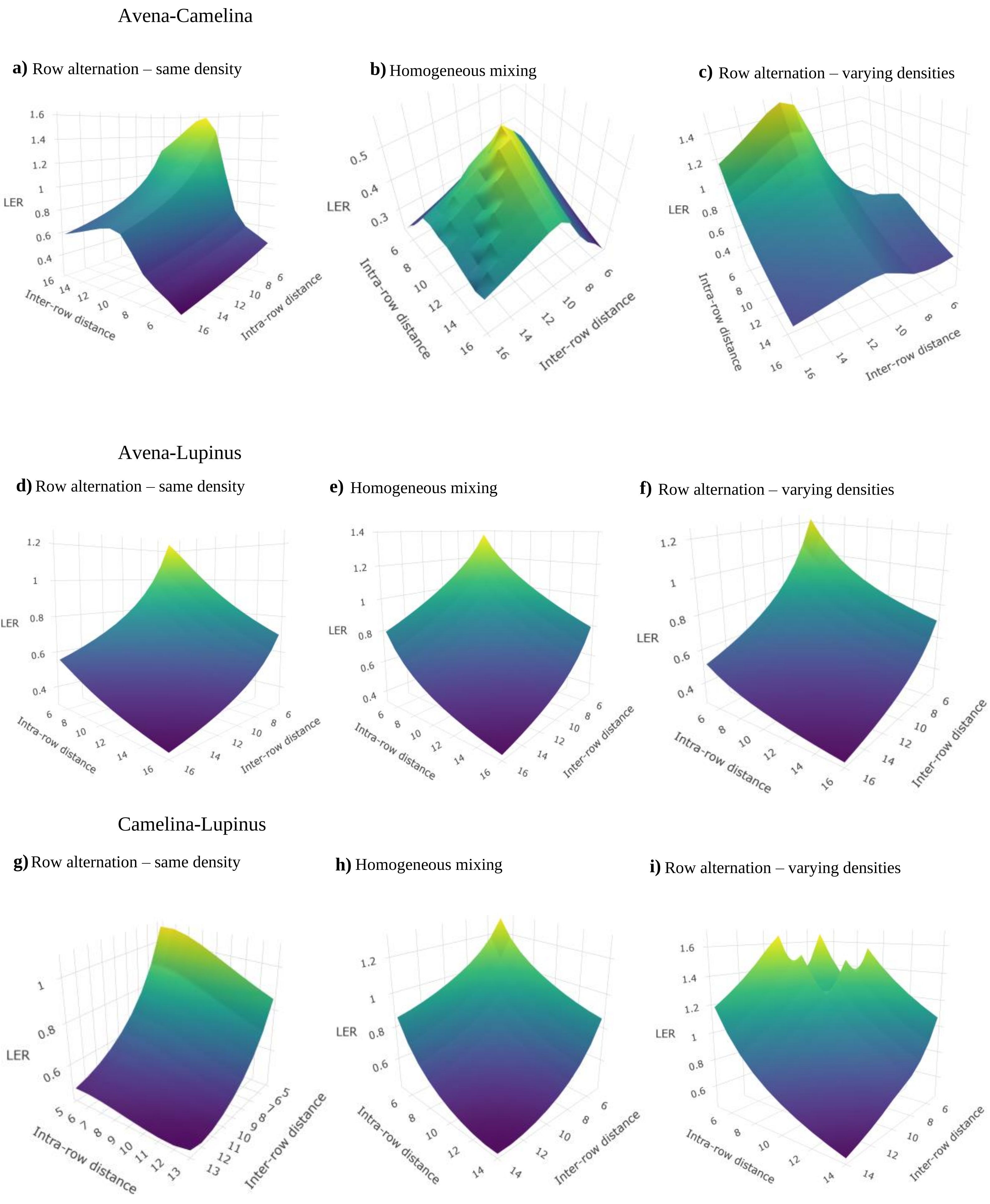


**Figure S2:** Surface plots showing LER in function of intra- and inter-row distance for two-species combinations, in various spatial configurations.

### Avena-Camelina-Lupinus

**a)** Row alternation – same density

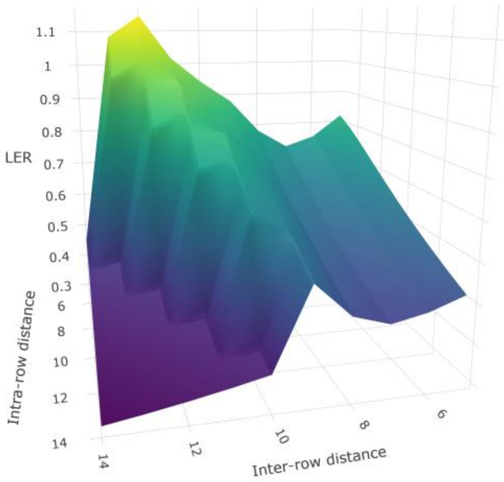


**b)** Homogeneous mixing

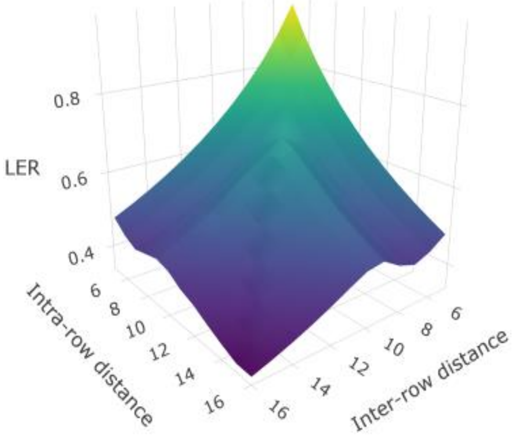


**c)** Random placement

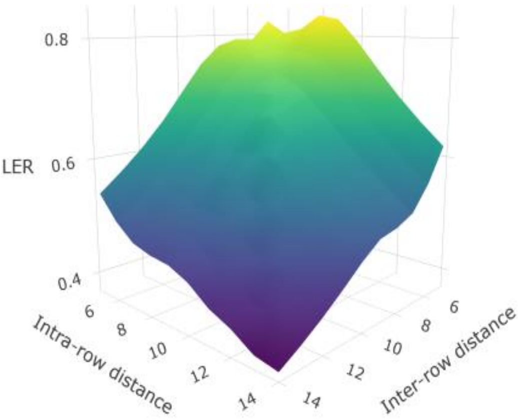


**d)** Row alternation – varying densities
(dens Ave = dens Lup = dens Cam/2)

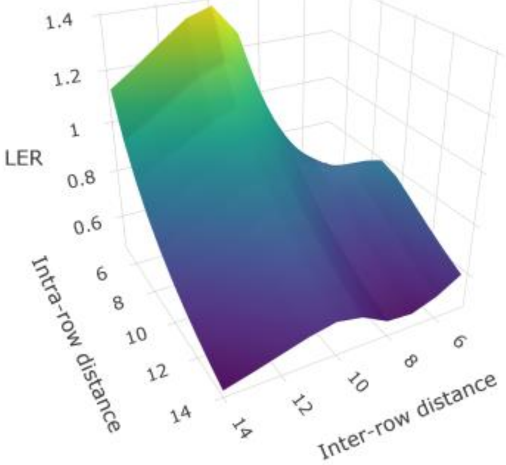


**e)** Row alternation – varying densities
(dens Lup = dens Cam/3, dens Ave = dens Cam/2)

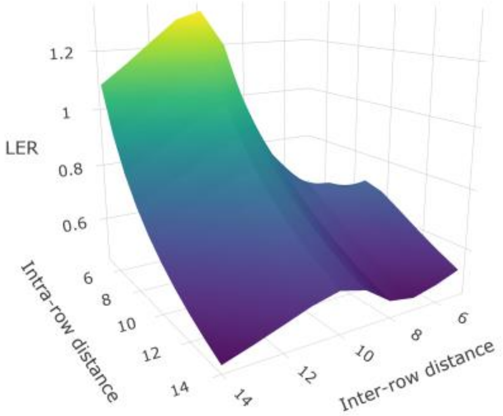


**Figure S3:** Surface plots showing LER in function of intra- and inter-row distance for the three species combinations Avena-Camelina-Lupinus, in various spatial configurations.

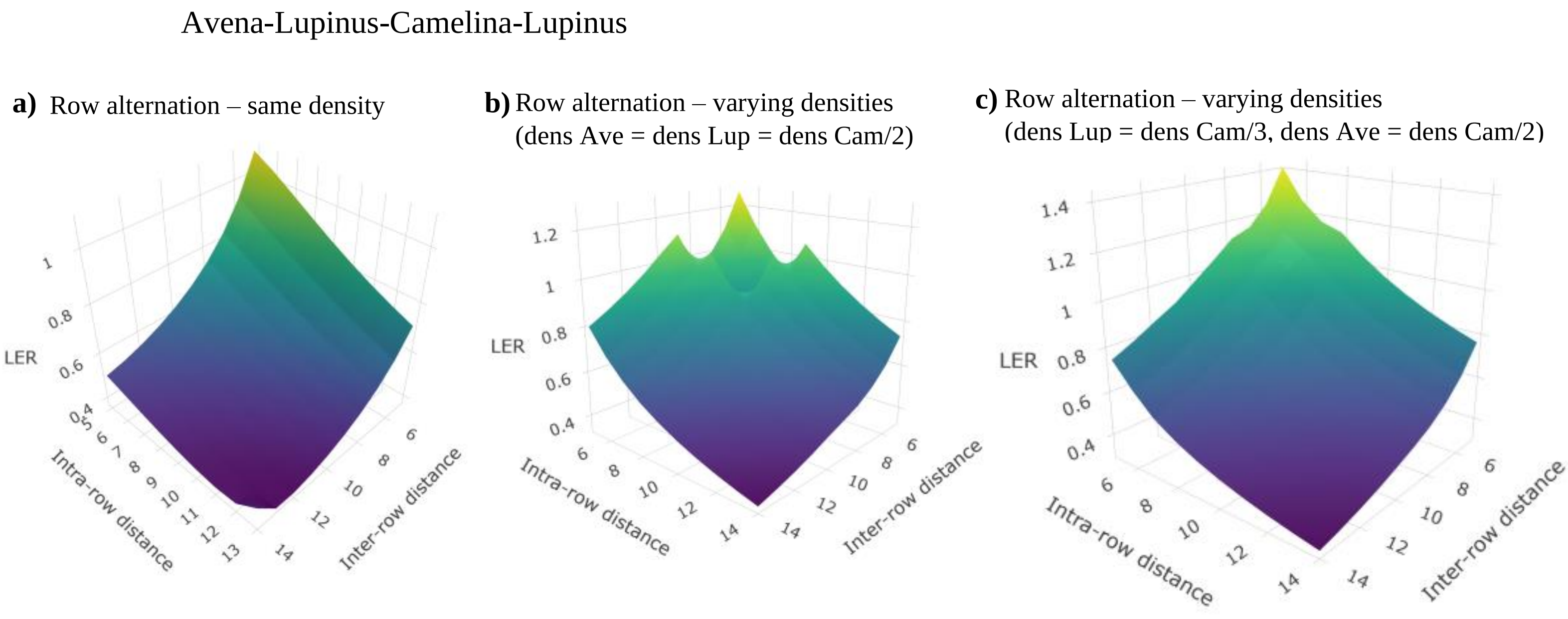


**Figure S4:** Surface plots showing LER in function of intra- and inter-row distance for the three species combinations Avena-Lupinus-Camelina-Lupinus, in various spatial configurations.
